# Parental transport induces a dormant state while maintaining oxytocin recruitment in poison frog tadpoles

**DOI:** 10.64898/2026.06.16.732608

**Authors:** Diogo F. Antunes, Zuyao Liu, Eva Ringler

## Abstract

Parental care can have pervasive effects on offspring’s neurodevelopment. Parent-offspring interactions are often modulated by the neuropeptide oxytocin, which is responsible for the development of social bonds. The development of the oxytocinergic system is dependent on the quality of parental care during the post-natal phase. However, it is yet unknown how post-natal direct interactions can influence the development of the oxytocinergic pathway. Here we tested how an obligate parental care behaviour, tadpole transport in poison frogs, influences the development of the oxytocinergic pathway. To this end, we quantified whole brain expression of oxytocin receptor and oxytocin precursor throughout three developmental stages of *A. femoralis* tadpoles, before, during and after tadpole transport. Our results show an overall downregulation during tadpole transport, which indicates that during transport tadpoles enter a dormant state to slow down development until they are placed in water. Interestingly, the expression of oxytocin precursor did not vary between the three developmental stages. This might indicate that oxytocin is being recruited during transport, but does not lead to neurodevelopmental changes. In sum, here we present the first evidence of a dormant state during tadpole transport which might be an adaptive response to the terrestrial reproduction in poison frogs.

## Introduction

Parental care comprises any behaviour performed by parents that increases offspring survival, typically at a cost to parental resources and future reproduction [1]. Despite the associated costs, parental care enhances lifetime reproductive success, leading to the evolution of parental care strategies in virtually all animal taxa (insects:[2]; teleosts: [3]; amphibians: [4]; birds and mammals: [5]). Although parental care strategies vary widely across the animal kingdom, many forms of parental care behaviours involve close physical contact between parents and offspring (e.g. licking and grooming and rats, mouth guarding in cichlids or brooding in frogs and salamanders). Consequently, there has been a great interest in how parent-offspring interactions shape offspring neurodevelopment and behaviour. For instance in rats, natural variation in maternal care alters the development of the hypothalamic-adrenal axis, leading to long-term changes in hippocampal glucocorticoid receptor expression and stress responsiveness [6]. Similarly, in the cooperatively breeding cichlid *Neolamprologus pulcher*, the presence or absence of parents during early development has lifelong consequences for dopamine and glucocorticoid receptor densities in several brain regions [7] and for baseline cortisol levels [8]. Despite growing evidence that parental care can have profound and lasting effects on offspring phenotype, the mechanisms by which direct parent-offspring interactions influence offspring neurodevelopment remain poorly understood.

Oxytocin is a well conserved neuropeptide across vertebrates [9], where it plays a central role in regulating social behaviour [10,11]. Recent work suggests that oxytocin contributes to emotional contagion and that this function is evolutionarily ancient [12]. In mammals, oxytocin is well known for mediating parent-offspring bounding and affiliative interactions [13], highlighting its importance for social regulation. The development of the oxytocinergic system itself can be shaped by parental care. In California mice, increased parental care leads to a higher number of oxytocin-expressing neurons in the paraventricular nucleus and the suprachiasmatic nucleus later in life [14]. Oxytocin is produced within a neuron expressing the *Oxytocin-Neurophysin 1* gene (*Oxt-Neu1*), whose peptide product is cleaved into oxytocin and Neurophysin-1 [15]. Hence, the quantification of OXT-Neu1 gene expression can serve as proxy for oxytocin synthesis and activity [16]. In combination with the expression of the oxytocin receptor (OxtR), this allows us to assess how parental care influences both ligand production and receptor recruitment during early postnatal development. However, it remains unknown how direct parent-offspring interactions during key developmental stages affect the development of the oxytocinergic system.

In several species of neotropical frogs, females deposit clutches in land, and after hatching tadpoles are transported by the parents to water bodies [17]. This phenomenon makes neotropical frogs the ideal model system to investigate the role of direct physical contact between parents and offspring for the development of the oxytocinergic pathway [17]. Particularly in the neotropical frog *Allobates femoralis*, tadpole transport behaviour has been intensively studied [18–22]. Males establish and defend territories, where females deposit clutches on land [23–25]. After hatching, males transport their tadpoles to small waterbodies in the forest [18], failure to transport may results in tadpole mortality either through desiccation or cannibalism by conspecific males [26,27]. In the absence of the father, females take over tadpole transport and compensate for the male’s absence, and ensure offspring survival [18,21]. During oviposition, females also deposit a gelatinous substance that helps to maintain the clutch hydrated and creates a semi-aquatic microenvironment [28,29], a trait that might have facilitated the evolutionary transition from aquatic to terrestrial breeding in amphibians. Because tadpole transport involved moving offspring from a semi-aquatic environment into open air and then into water, it can potentially be a critical transitional phase in the tadpole’s development. During this period, offspring experience direct physical contact with the transporting parent and get exposed to a novel medium, both of which may influence neurodevelopment. Given the dire consequences of transport failure, tadpole transport can be a critical phase for the parentally-induced development of the oxytocinergic system. We therefore hypothesize that parental transport will later *Oxt-Neu1* expression and subsequently modulate *OxtR* expression upon deposition in water, reflecting coordinated changes in the oxytocin synthesis and receptor recruitment during early development. Additionally, we also predict that the transient exposure to air during transport could induce transcriptional changes associated with the shift from a semi-aquatic to an aerial environment.

## Methods

### Subjects and housing conditions

All tadpoles used in the current study originated from a *Allobates femoralis* population that is kept and breed at the animal facilities of the Hasli Ethological station of the University of Bern. Adult *A. femoralis* are housed in pairs in terrariums of 60x40x40cm and 50x40x40cm. All animals are kept under a light: dark cycle of 12:12h, temperature ranging from 21±1°C to 28±1°C and rain twice a day for 5min. The housing conditions mimic the natural conditions in French Guiana [30]. All terrariums are equipped with expanded clay pebbles and dry oak leaves on the floor, half a coconut shell as a shelter, a small tree log for perching, a water bowl, and plants for enrichment. All terrariums are partially covered with tree fern stems, cork mats, and a curtain in the front to reduce visual contact between neighboring terrariums. Adults are fed with wingless fruit flies dusted in vitamin powder (Amphib, Herpetal) twice a week.

### Allobates Femoralis coding sequences

We started by identifying *A. femoralis*’ orthologue sequences for Oxytocin receptor (OxtR), Oxytocin-Neurophysin-1 (Oxt-Neu1), 18S ribosomal RNA and glyceraldehysde-3-phosphate dehydrogenase (GAPDH). We used publicly available protein sequences (see Table 1) to run a Basic Local Alignment Search Tool (tBLASTn, NCBI [31]) against *A. femoralis’* genome (NCBI genome assembly GCA_033576535.1) to identify the candidate scaffold containing our gene of interest. We then extracted the candidate scaffold from *A. femoralis* genome and ran Exonerate [32] using the protein2genome model to predict the structure of our gene of interest (i.e the position of the introns and exons). Using the predicted coordinates of the exons, we extracted the coding sequence and ran a BLAST in NCBI to validate our targets. After the validation, we designed primers spanning exon-exon junctions using Primer3 [33]. Primers were then tested by touch-down PCR and amplicons were sequenced to validate primer specificity.

**Table 1:**
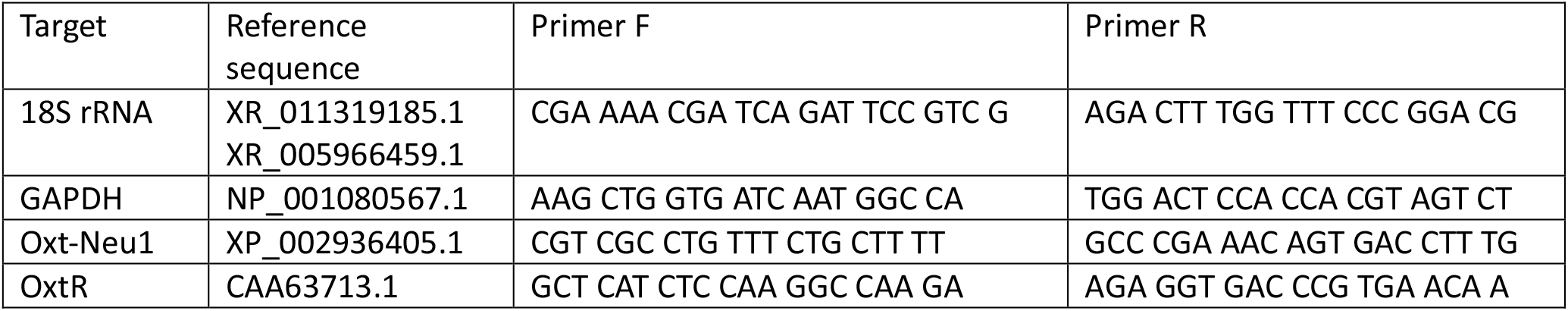
Reference sequences used to for the tBLASTn against *A. femoralis* genome and primers used in RT-qPCR experiment designed using Primer3. For 18S rRNA used two reference sequences, one from *Ranitomeya imitator* and one sequence from *Xenopus laevis*, and sequences identified the same target in *A. femoralis*.

### Tissue collection, RNA extraction

Ten tadpoles per developmental stage (before, during and after male transport) were collected (N=40) and were euthanized with an overdose of buffered MS222 (MERCK). The whole brains were quickly dissected under a dissection microscope (WILD M3C) and placed in 150µL of DNA/RNA shield (Zymo Research) and stored at -80°C until further processing. Prior to total RNA extraction, whole brains were digested with proteinase K at 55°C for 1h 30min. Total RNA extraction was performed using quick RNA mini-prep (Zymo Research) following the manufacturer’s protocol, including an in-column DNAse treatment to avoid genomic contamination. To increase RNA concentration the total elution volume was reduced to 50µL. Total RNA was quantified using QuBit HS-RNA kit (Thermo-Fischer), following the manufacturer’s protocol. cDNA conversion was performed using iSCRIPT (Bio-Rad), standardizing RNA input to 350 ng across all samples.

### Quantitative real-time PCR

To determine the amplification efficiency (E) and the absence of primer dimers for all primers, quantitative PCR experiments and amplicon melting curves (50-90°C) were run using standard curves of 4x5-fold dilutions of all brain RNA in triplicate [7,34]. RT-qPCR experiments were prepared in 96-well plate (Thermo-Fischer) by combining our primers (Microsynth), 1uL of cDNA (from a concentration of 17.3ng/µL) and 5x HOT FIREPol EvaGreen qPCR Mix (Solis BioDyne) on an Aria-Mx (Agilent). All samples were run in triplicate with no-template controls. To verify that only a single-amplified product was produced and to confirm the absence of primer dimers, melting curves were performed for each replicate. Cycle thresholds (Ct) for each sample were used to calculate gene expression for each individual following the formula E^-Ct^.

### Statistics

All statistical analyses were conducted using the software R version 4.3.3. Some samples were excluded from the analysis when the coefficient of variance (CV) of the three replicates was larger than 5%. We started by testing the pairwise expression stability of our candidate housekeeping genes 18s and GAPDH using “geNorm” [35]. Despite the M value being within the acceptable expression stability (M=0.72), this does not mean that the expression of 18s and GAPDH do not vary across biological stages. To test for their biological stability, we fitted linear mixed-effects models (LMM) using “lme4” and “lmerTest” [36,37]. LMMs were fitted with the log_2_-transformed expression as response variable, developmental stage (before, during or after transport) as fixed effect and parent ID as a random effect. Model assumptions were tested using the package DHARMa. When the random factor in our models did not explain any variation, the random term was removed and linear models were fitted (Bates, https://stat.ethz.ch/pipermail/r--sig--mixed--models/2014q3/022509.html). Outliers were identified and removed by calculating the Cook’s distance of the model’s residuals using the function “cooks.distance” from the package “stats”. Due to the differential expression of both candidate housekeeping genes (see Results section) showing that their expression has biological meaning, hence we cannot use either genes as a control for the expression of our genes of interest. For this reason, all our analysis reflects the expression level in relation to the standardized cDNA template for every RT-qPCR reaction (17.5 ng of cDNA). The effect of developmental stage on the expression of oxytocin receptor and oxytocin-neurophysin 1 were analyzed by fitting LMM, following the same procedure described above. Estimated marginal means (EMMs) for each developmental stage were calculated from the models using the package “emmeans”, and pairwise comparisons were corrected for multiple testing following Tukey’s method. All visualizations were generated using the package “ggplot2”.

## Results

The expression of our candidate housekeeping genes was significantly different across developmental stages (Table 2). For *18S*, expression during transport was significantly lower than after transport, while there were no significant differences between the other developmental stages (Table 3, Fig. 1a, Fig. 1b). *GADPH* showed a similar expression pattern, with significantly lower expression during transport when compared to after transport, and there were no expression differences between before and after transport.

**Table 2:**
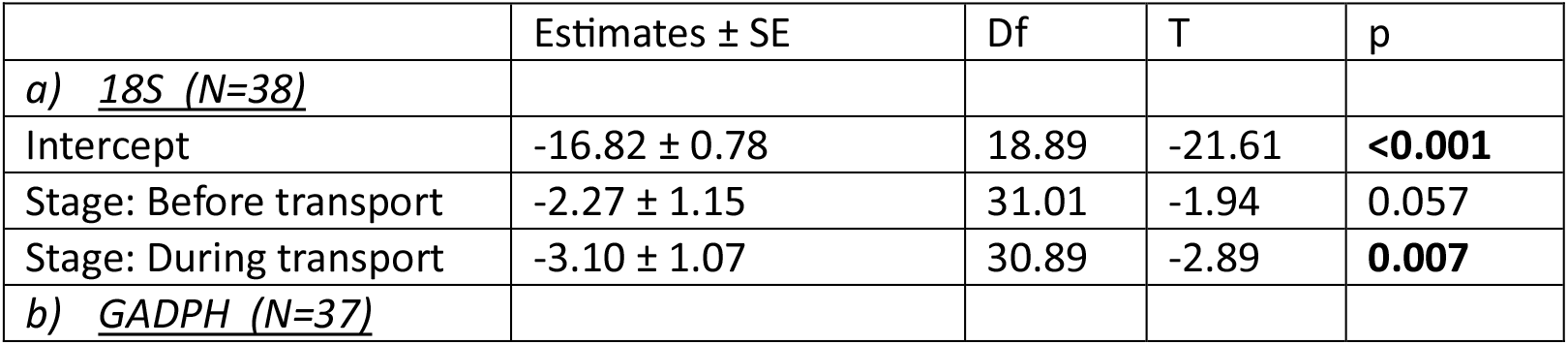
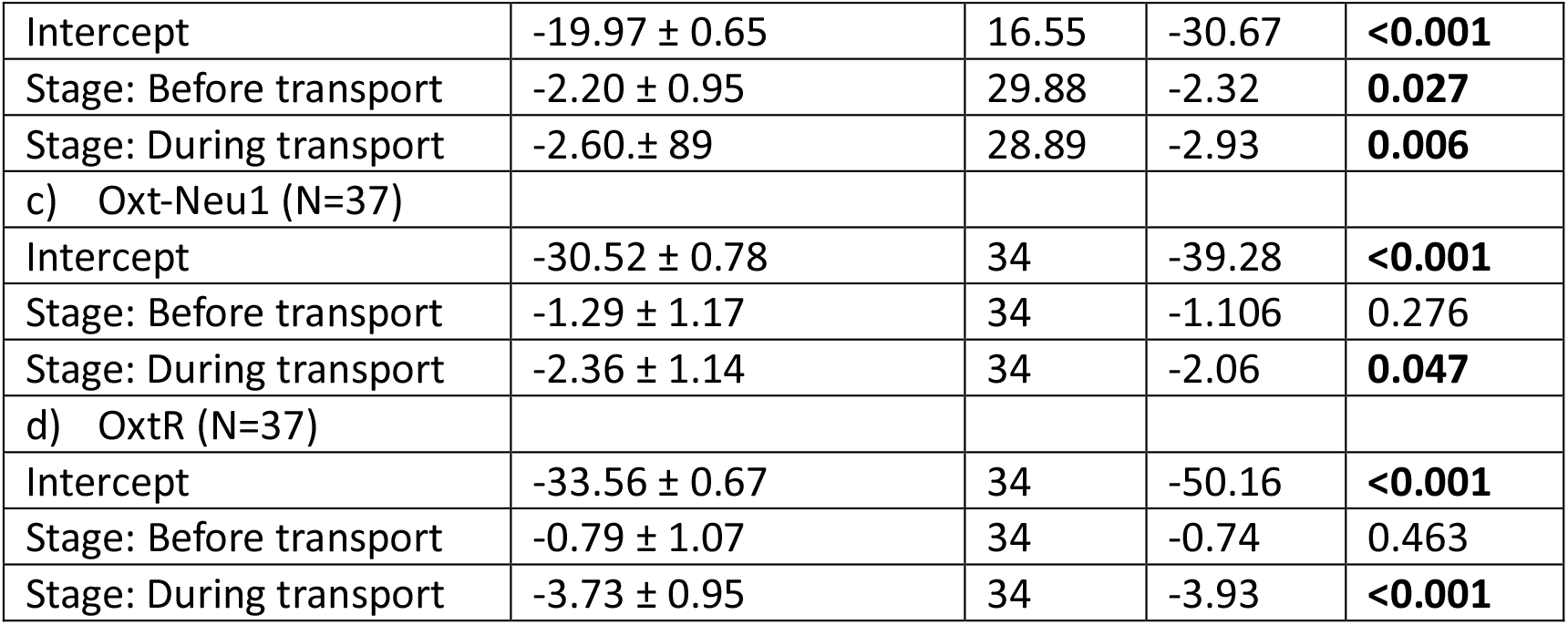
Results from the LMM on the effect of tadpole developmental stage on the expression of (a) *18S* rRNA, (b) *GADPH*, (c) *Oxt-Neu1* and (d) *OxtR*.

| | Estimates ± SE | Df | T | p |
| --- | --- | --- | --- | --- |
| **a) 18S (N=38)** |  |  |  |  |
| Intercept | -16.82 ± 0.78 | 18.89 | -21.61 | **<0.001** |
| Stage: Before transport | -2.27 ± 1.15 | 31.01 | -1.94 | 0.057 |
| Stage: During transport | -3.10 ± 1.07 | 30.89 | -2.89 | **0.007** |
| **b) GAPDH (N=37)** |  |  |  |  |
| Intercept | -19.97 ± 0.65 | 16.55 | -30.67 | **<0.001** |
| Stage: Before transport | -2.20 ± 0.95 | 29.88 | -2.32 | **0.027** |
| Stage: During transport | -2.60 ± 0.89 | 28.89 | -2.93 | **0.006** |
| c) *Oxt-Neu1* (N=37) |  |  |  |  |
| Intercept | -30.52 ± 0.78 | 34 | -39.28 | **<0.001** |
| Stage: Before transport | -1.29 ± 1.17 | 34 | -1.106 | 0.276 |
| Stage: During transport | -2.36 ± 1.14 | 34 | -2.06 | **0.047** |
| d) *OxtR* (N=37) |  |  |  |  |
| Intercept | -33.56 ± 0.67 | 34 | -50.16 | **<0.001** |
| Stage: Before transport | -0.79 ± 1.07 | 34 | -0.74 | 0.463 |
| Stage: During transport | -3.73 ± 0.95 | 34 | -3.93 | **<0.001** |

**Table 3:** Results from the pair-wise comparisons followed by Tukey multiple testing correction on the expression of (a) *18S* rRNA, (b) *GAPDH*, (c) *Oxt-Neu1* and (d) *OxtR*.

|  | Estimates ± SE | Df | T ratio | p |
| --- | --- | --- | --- | --- |
| a) *18S* (N=38) |  |  |  |  |
| After Vs before transport | 2.27 ± 1.21 | 31 | 1.88 | 0.163 |
| After Vs during transport | 3.105 ± 1.09 | 30.9 | 2.84 | **0.021** |
| Before Vs during transport | 0.832 ± 1.22 | 32.1 | 0.68 | 0.776 |
| b) *GADPH* (N=37) |  |  |  |  |
| After Vs before transport | 2.202 ± 0.999 | 30.4 | 2.204 | 0.086 |
| After Vs During transport | 2.597 ± 0.905 | 29.5 | 2.869 | **0.02** |
| Before Vs During transport | 0.395 ± 1.020 | 32.3 | 0.387 | 0.921 |
| c) *Oxt-Neu1* (N=37) |  |  |  |  |
| After Vs before transport | 1.30 ± 1.17 | 34 | 1.11 | 0.517 |
| After Vs During transport | 2.36 ± 1.14 | 34 | 2.064 | 0.113 |
| Before Vs During transport | 1.06 ± 1.21 | 34 | 0.877 | 0.658 |
| d) *OxtR* (N=37) |  |  |  |  |
| After Vs before transport | 0.79 ± 1.07 | 34 | 0.742 | 0.741 |
| After Vs During transport | 3.72 ± 0.95 | 34 | 3.93 | **0.001** |
| Before Vs During transport | 2.93 ± 1.07 | 34 | 2.74 | **0.026** |

**Figure 1:**
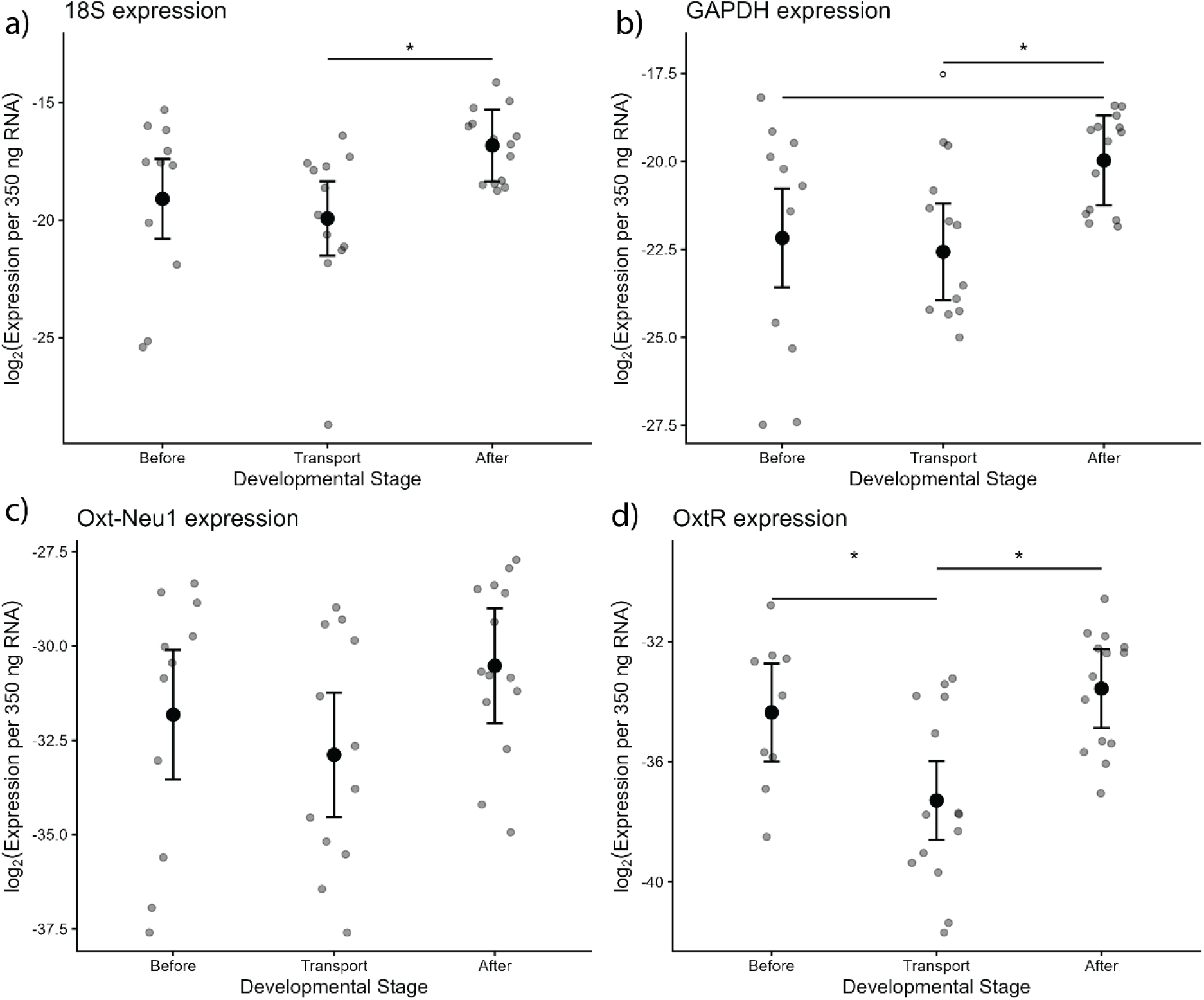
Model estimated expression across developmental stages for a) 18S rRNA, b) GAPDH, c) Oxt-Neu1 and d) OxtR. Gray points represent raw log2-transformed expression values per tadpole. Solid black points represent the predicted means from the LMM. Bars represent confidence intervals. Significant effects from pair-wise comparisons are represented with an asterisk. Statistical trends are represented by an unfilled circle.

For our target genes, *OxtR* expression was significantly different across developmental stages (Table 2). Its expression was significantly lower during transport when compared to both before and after transport (Table 3, Fig. 1d). In contrast, the expression of *Oxt-Neu1* did not differ across developmental stages (Table 3 and 4, Fig.1c).

## Discussion

Here we investigate whether parental tadpole transport influences the development of the oxytocinergic system in *Allobates femoralis*. Contrary to our initial prediction, we found no evidence that parental transport increases *Oxt-Neu1* expression, indicating that oxytocin synthesis remains stable across developmental stages. In contrast, *OxtR* expression is significantly lower during tadpole transport, when compared to before and after transport. We found a similar reduction in the expression of *18S* and *GAPDH*, both of which are associated with cellular metabolism. The observed coordinated decrease in transcription suggests that tadpoles may transiently reduce transcriptional activity during transport, potentially reducing their metabolic state and enter a dormant state to cope with the transition from semi-aquatic to aerial environment.

Contrary to our initial prediction, the development of the oxytocinergic pathway did not depend on tadpole transport. We show that the oxytocin precursor did not differ across developmental stages, indicating that oxytocin and neurophysin-1 synthesis remains stable throughout development. Interestingly, Oxt-Neu1 was the only transcript studied here, that did not differ across stages. Based on these results, we hypothesize that paternal tadpole transport sustains oxytocinergic activity, reflected in the stable recruitment of the oxytocin precursor despite the overall transcription reduction. The effect of parental care on the development of the oxytocinergic system is a well a known phenomenon. For example, female rats raised with levels of maternal licking and grooming show an increase in oxytocin receptor density in the amygdala and bed nucleus of the stria terminalis [38]. In California mice, offspring that receive high levels of care have a higher number of oxytocin positive cells in the paraventricular nucleus and supraoptic nucleus [14]. Parental care can also influence circulating oxytocin levels, children receiving higher maternal care have higher plasma oxytocin levels [39]. In our study, tadpoles maintain a stable expression of oxytocin precursor (Table 3; Fig 1c), raising the hypothesis that physical contact with the father during transport may induce oxytocinergic activity. However, manipulations of tadpole transport are needed to test this hypothesis. It remains yet unknown whether female tadpole transport compensation leads to similar effects on oxytocinergic activity [21].

Our study revealed an unexpected reduction in the expression of *18S, GAPDH* and *OxtR* during tadpole transport, with expression levels returning to pre-transport levels after being deposited in water. The decrease in transcripts associated with cellular function was particularly surprising. The *18S* ribosomal RNA encodes a key component of the small ribosomal subunit and is essential for protein synthesis [40,41]. Reduced *18S* levels have been shown to impair translation and slow cell growth in mammalian cells [42], and cellular stress can trigger 18S degradation [41]. *GAPDH*, a central enzyme in ATP production and glycolysis [43], is ubiquitously expressed across taxa [44], and reduced GAPDH expression can limit cell proliferation [45]. Considering the functions of both genes, the coordinated lower expression observed during transport suggests that *A. femoralis* tadpoles may temporarily reduce metabolic activity and development, entering a transient dormant state during transport. Similar broad transcriptional down-regulation has been described in hibernating vertebrates. In golden-mantled squirrels exhibit reduced transcriptional activity during torpor [46]. We hypothesize that the reduced transcriptional activity observed during transport might be a response to the transition from a semi-aquatic environment to the exposure to air. Exposure to air prior to metamorphosis may increase oxidative stress due to the higher oxygen availability [29]. Entering a dormant state during transport could therefore help mitigate potential developmental costs associated with oxidative stress. Further studies are needed to validate this hypothesis.

In conclusion, we show that expression of oxytocin precursor remains stable across developmental stages, despite the overall reduction of transcriptional activity during tadpole transport. We also show a general decrease in transcriptional activity during transport, which is restored once the tadpoles are deposited in water. To our knowledge, these results provide the first evidence of reduced cellular activity during tadpole transport in Neotropical frogs. We hypothesize that tadpoles may enter a transient dormant state during transport to mitigate potential physiological challenges associated with the exposure to air prior to metamorphosis. Such a response, together with the evolution of parental care and the emergence of water-retaining clutch jelly, may have facilitated the transition from aquatic to terrestrial oviposition in neotropical poison frogs.

## Acknowledgements

We are grateful for the logistic support provided by Lauriane Bégué and Evi Zwygart. We thank Marius Rösti for his insights throughout the project, Maddy Thakur and Katie Peichel for providing the necessary laboratorial equipment for the execution of this project. We acknowledge the financial support by the Swiss National Science foundation (SNSF, project 310030_215049) to E.R.

